# Dynamical Systems Modeling of Bt Resistant Cornborer Population Dynamics

**DOI:** 10.1101/795682

**Authors:** Jomar F. Rabajante, Arian J. Jacildo, Edwin P. Alcantara

## Abstract

Mathematical models provide insights for the design and optimization of strategies to control disease epidemics and evolution of pesticide resistance in crops. Here, we present a simple mathematical model to investigate the population dynamics of non-Bt resistant and Bt resistant cornborers in a field with refuge. In the presence of refuge, it is expected that the population of Bt resistant pests will decline due to the dilution effect (non-Bt resistant cornborers mate with Bt resistant pests). We have found that increasing the refuge size can be effective in reducing Bt resistant pests as long as there exists a relatively huge population of non-Bt resistant cornborers at the start of the simulation. This implies that refuge is useless in inhibiting the evolution of cornborers if a sufficient initial population of non-Bt resistant cornborers is absent.

## Main Text

Mathematical models are useful tools in predicting the incidence and prevalence of plant pests and diseases [Madded et al., 2007; Donatelli et al., 2017]. Analysis of the mathematical models provide essential insights for the design and optimization of strategies to control disease epidemics and evolution of pesticide resistance [Tyutyunov et al., 2007; Vyska et al., 2016; Cortez et al., 2017]. In maize crops, various models have been formulated to investigate the epidemiology and spread of cornborers [Tyutyunov et al., 2007; Sudo et al., 2017]. There are models that investigated the efficacy of high-dose refuge strategy which can prevent or delay the occurrence of Bt resistant pests [Tyutyunov et al., 2007; Campagne et al., 2016]. Here, we present an example of a simple mathematical model that we have designed to study the population dynamics of non-Bt resistant and Bt resistant cornborers in a field with refuge. Furthermore, we discuss here several implications of the model results that can be potential strategies for controlling the incidence and prevalence of Bt resistant pests.

We model the population dynamics of cornborers using differential equations, which is commonly used in investigating systems that are temporally dynamic. The spatial characteristics of the field are simplified and incorporated in the *K*_*refuge*_ and *K*_*Bt*_ parameters that respectively represent the relative size of the refuge and Bt corn area, and in the parameter *γ* that represents spatial dispersion of cornborers. The definitions of the state variables and parameters in the model are presented in Table 1. The change in the population frequency of non-Bt resistant cornborer per unit time is modeled by

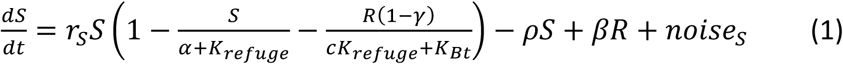

and the change in the population frequency of Bt resistant cornborer is represented by

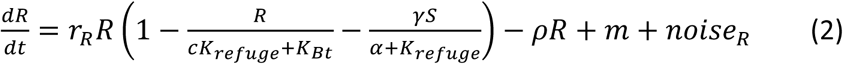

**Table 1.** Definition of the state variables and parameters. Unless otherwise stated, the simulations shown in the figures use the default values presented in this table [Campagne et al. 2016; Tyutyunov et al., 2007; Song et al., 2013; Sudo et al., 2017].

| Parameter | Definition | Default value |
| --- | --- | --- |
| $S$ | population frequency of non-Bt resistant cornborers | Variable |
| $R$ | population frequency of Bt resistant cornborers | Variable |
| $r_S$ | Reproduction coefficient of non-Bt resistant cornborer | 150 |
| $r_R$ | Reproduction coefficient of Bt resistant cornborer | 150 |
| $K_{Refuge}$ | Relative size of the refuge measured in terms of the carrying capacity for the cornborers | 0.1 |
| $K_{Bt}$ | Relative size of the Bt corn area measured in terms of the carrying capacity for the cornborers | 0.9 |
| $1 - c$ | Fitness cost of Bt resistant cornborer when foraging at the refuge | 0.1 |
| $\alpha$ | Additional survival area for adult non-Bt resistant cornborers | 0.1 |
| $1 - \gamma$ | Probability of randomly acquiring a Bt resistant gene through mating and spatial dispersion of cornborers | 0.75 |
| $\rho$ | Natural or controlled death rate of cornborers | 0.75 |
| $\beta$ | Stock up rate for added population of non-Bt resistant cornborer | 0.1 |
| $\lambda$ | Average random occurrence of Bt cornborer mutants per unit time | 0.001 |
| $\sigma$ | Amplitude of environmental noise | 0.1 |

where

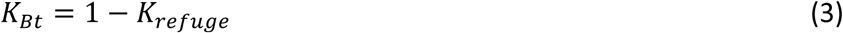

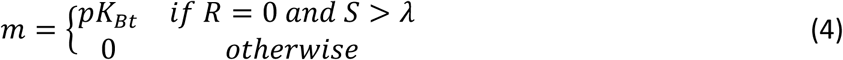

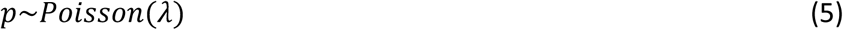

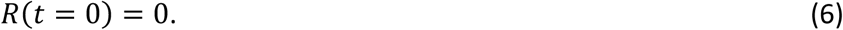

Equations 1 and 2 are modeled using the assumption that the growth rate of the cornborer population follows a logistic curve [Madded et al, 2007; Tyutyunov et al., 2007]. The carrying capacity of the non-Bt resistant cornborer population is determined by the relative size of the refuge plus other areas where adults can survive and reproduce. On the other hand, the carrying capacity of the Bt resistant cornborer population is capped by the size of the Bt corn area and refuge subject to fitness cost. The populations of the non-Bt resistant and Bt resistant cornborer are subject to evolutionary selection, which is assumed to be represented by the competition term in the logistic function. The model also includes mortality rate with *ρ* as the parameter, which can represent death due to natural means or through non-Bt control strategies (e.g., pesticides). Furthermore, there is an option to introduce a new set of non-Bt resistant cornborers through time to help dilute the population of Bt resistant pests. This strategy is assumed to be dependent on the current frequency of Bt resistant pests with stock up rate equal to *β*.

It is assumed in the simulations that the initial population frequency of Bt resistant cornborers is zero. The occurrence of mutants follows a Poisson distribution with mean *λ* and proportional to the size of the Bt corn area. Moreover, environmental noise, which represents unaccounted random events, affects the dynamics of the cornborer populations. The noise terms are modeled by the following:

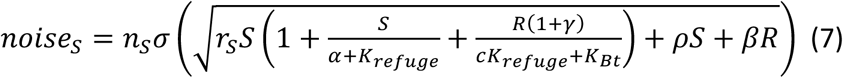

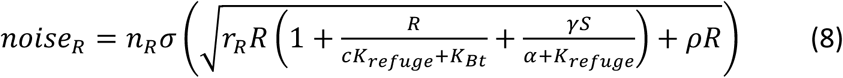

where *n*_*s*_, *n*_*R*_ ∼ *Normal*(0,1).

In our simulations, the population frequency of Bt resistant cornborers at a certain period of time can be predicted. Figure 1 shows an example of population dynamics based on one simulation run. In this figure, the Bt resistant population increases rapidly when the population frequency reaches around 5%. Since the model is probabilistic (e.g., occurrence of a mutant is random), one simulation run is not enough to visualize the different scenarios that can happen. Figures 2 and 3 present model results based on 1000 simulation runs. The simulation runs are summarized using the mean of the population frequencies (here, termed as the average case), and mean plus standard deviation (worst case).

**Figure 1.**
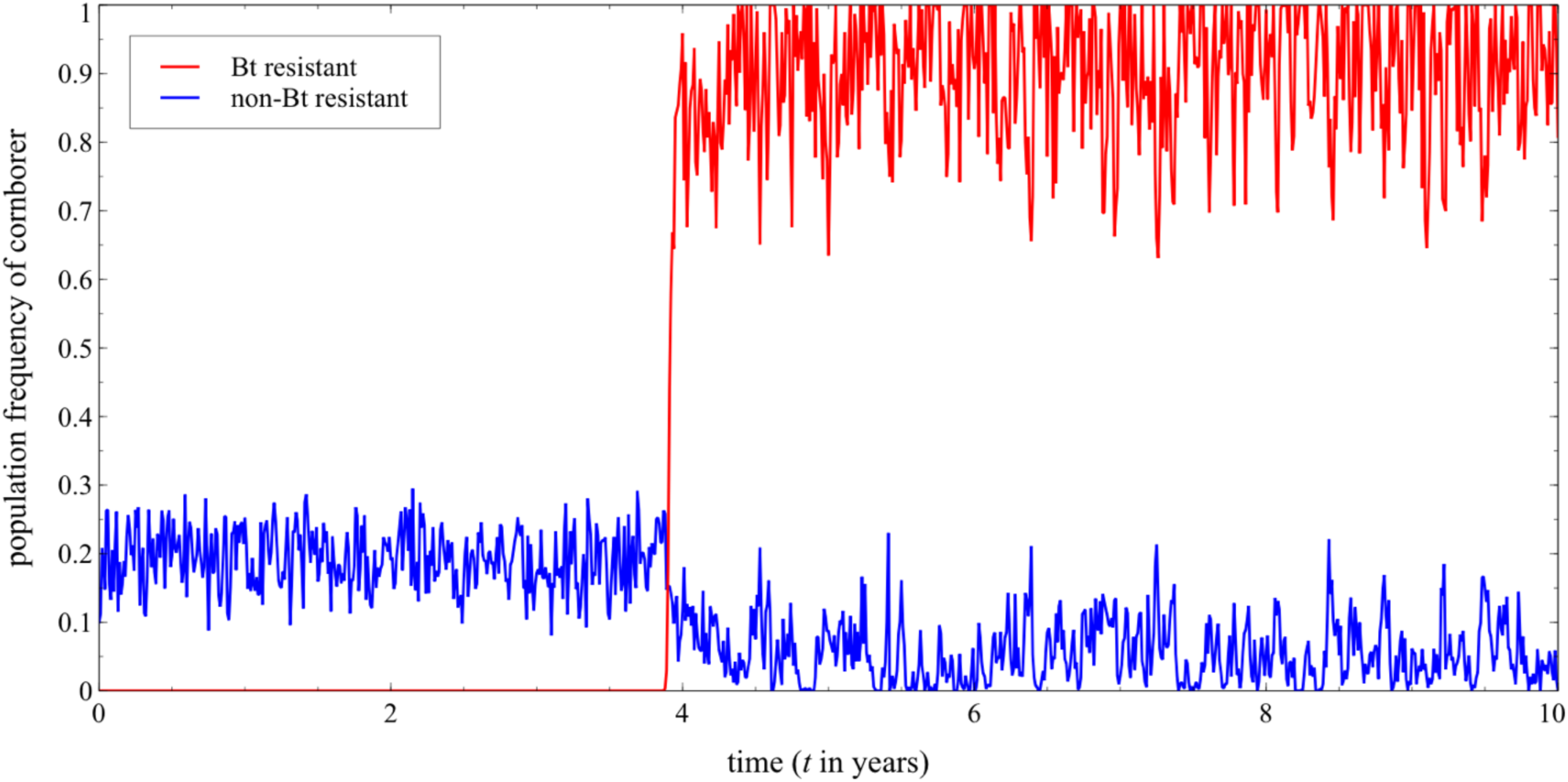
Sample simulation that shows Bt resistant cornborers eventually become dominant in around 3-4 years. The Bt resistant population increases rapidly when its population frequency reaches around 5%.

**Figure 2.**
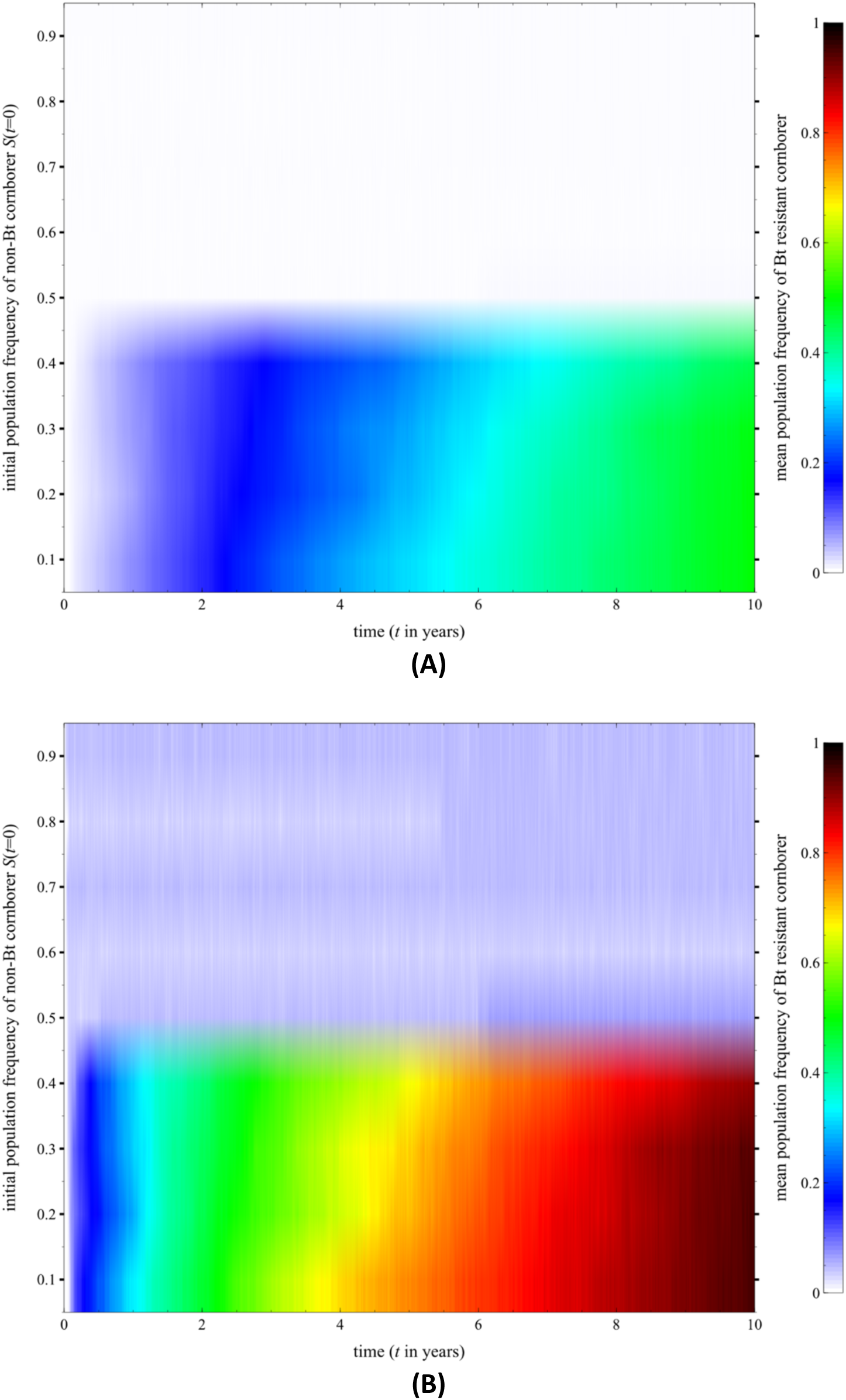
Initial population frequency of the non-Bt resistant cornborers (*S*(*t* = 0)) affects the persistence of Bt resistant cornborers. Results are based on the mean of 1000 simulation runs. **(A)** Average Case. **(B)** Worst Case (with standard deviation).

**Figure 3.**
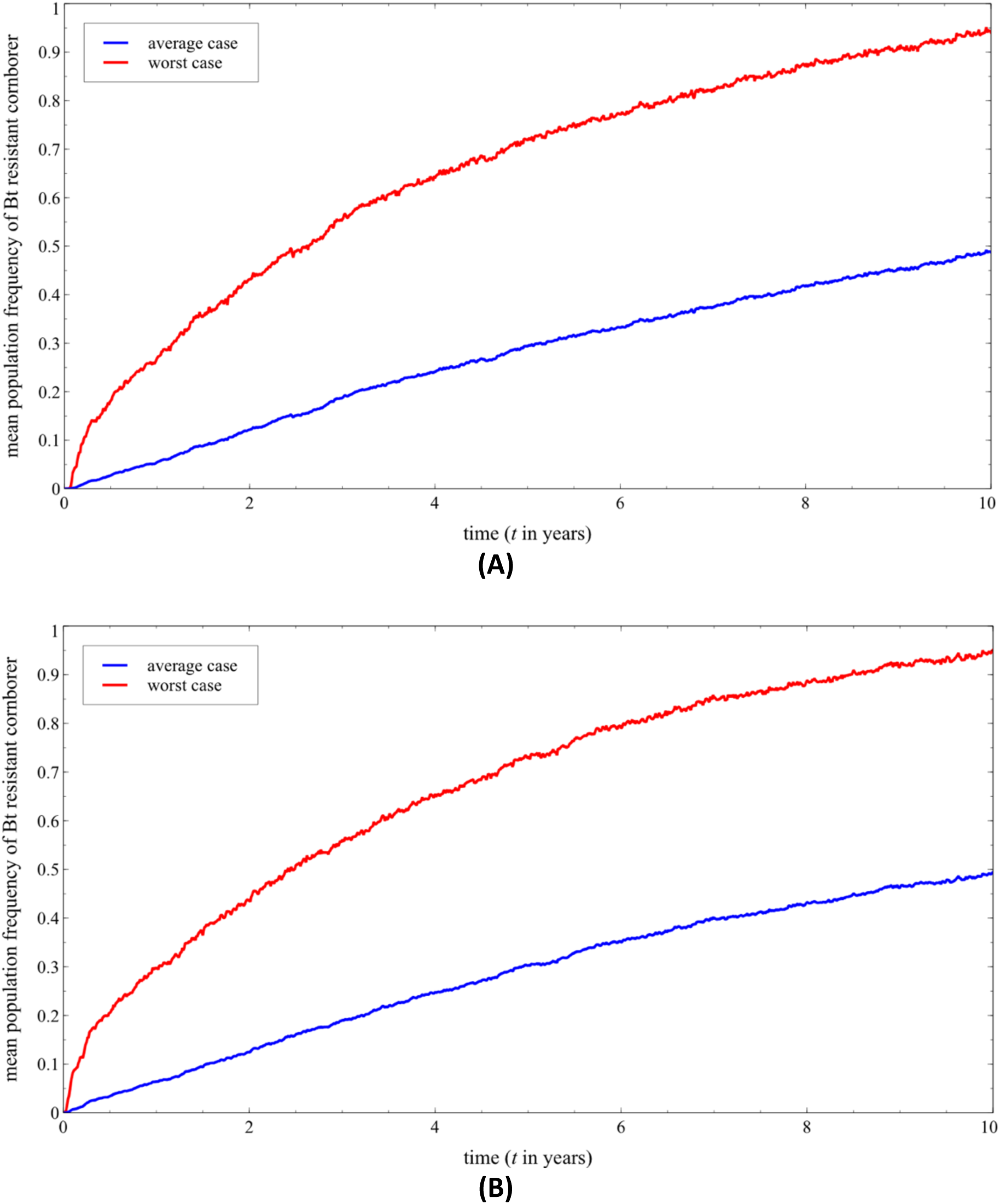
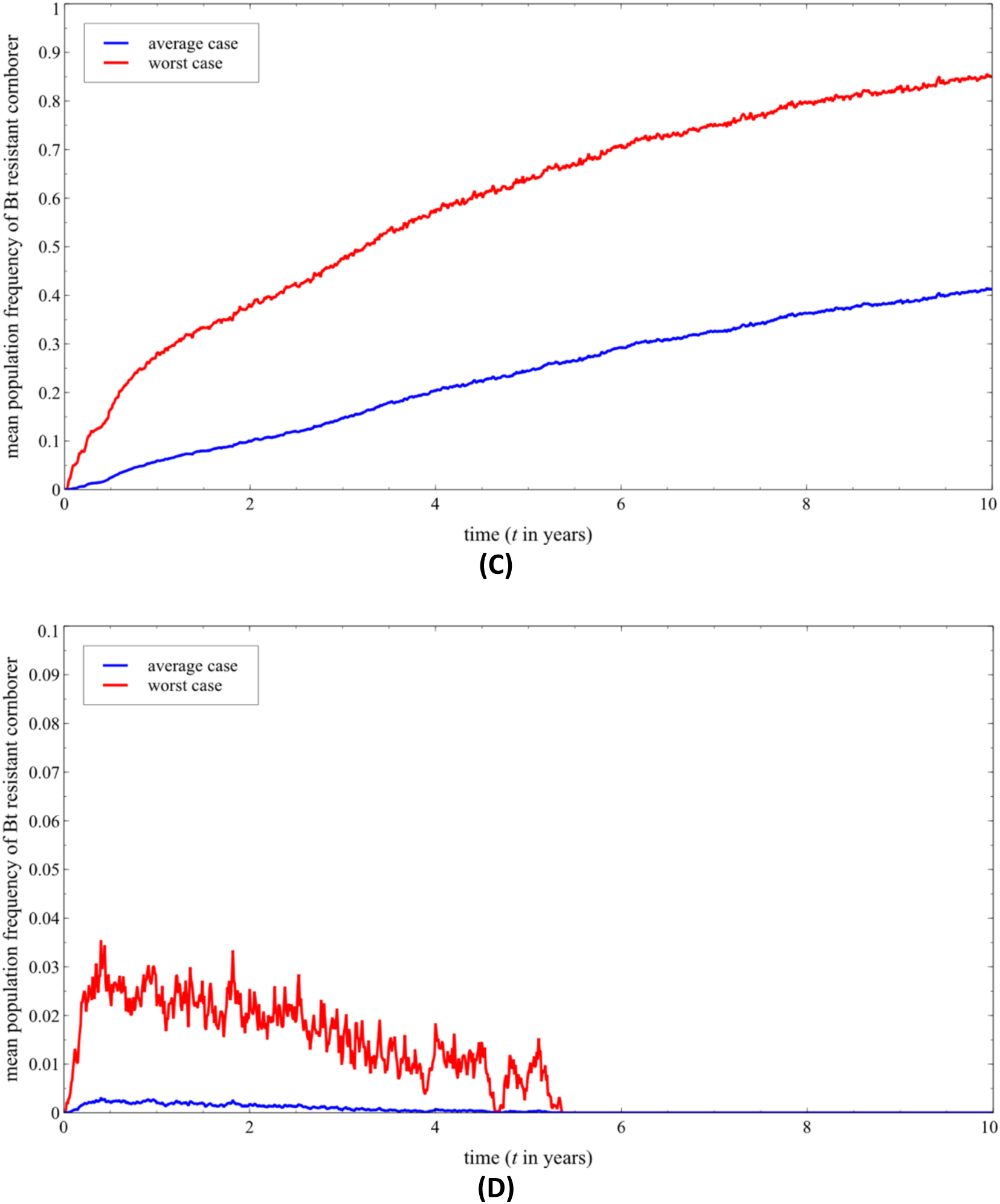
Effect of refuge size and mortality rate of cornborers to the population dynamics of Bt resistant pests. **(A)** *K*_*Refuge*_ = 0.05. **(B)** *K*_*Refuge*_ = 0.1. **(C)** *K*_*Refuge*_ = 0.5. For Figures 3A-3C, *ρ* = 0.75. **(D)** *ρ* = 112.5, which includes removal of eggs. Refuge size has a minimal effect on the persistence of Bt resistant cornborers, while mortality (e.g., due to pesticide application or biocontrol) helps in eliminating Bt resistant pests.

The model predicts that the initial population frequency of the non-Bt resistant cornborers (*S*(*t* = 0)) affects the persistence of Bt resistant cornborers (Figures 2). If *S*(*t* = 0) is low (e.g., <50%), it is more likely that the Bt resistant pests will persist. Figure 2A and 2B show the average and worst cases where population frequency of Bt resistant cornborers reaches around 60% and nearly 100% in ten years, respectively. On the other hand, mutant Bt resistant cornborer population is more likely not to be selected when a large initial population of non-Bt resistant cornborers is present (e.g., *S*(*t* = 0) > 50%). This may be due to the dilution effect, where frequency of Bt resistant genes declines as non-Bt resistant cornborers mate with Bt resistant pests.

Figures 3A-3C show the effect of refuge size on the population dynamics of Bt resistant pests. In the absence of refuge, it is expected that Bt resistant pests persist. Given a 10% initial population frequency of non-Bt resistant cornborers, a refuge size *K*_*Refuge*_ = 0.05 has similar result as *K*_*Refuge*_ = 0.1 (Figure 3A and 3B). A larger refuge size *K*_*Refuge*_ = 0.5 results in a small improvement (Figure 3C) but not as significant if we increase the initial population frequency of non-Bt resistant cornborers to 50% (Figure 2). This implies that increasing the refuge size can be effective in reducing Bt resistant pests as long as there exists a relatively huge population of non-Bt resistant cornborers. In addition, mortality (e.g., due to pesticide application, removal of insect eggs, and biocontrol) has huge impact in eliminating Bt resistant pests as expected (Figure 3D). This can be further studied by extending the model to include factors that can increase the death rate of cornborers.

